# Allosteric modulation of the Lon protease by effector binding and local charges

**DOI:** 10.1101/2024.09.06.611642

**Authors:** Justyne L Ogdahl, Peter Chien

## Abstract

The ATPase Associated with diverse cellular Activities (AAA+) family of proteases play crucial roles in cellular proteolysis and stress responses. Like other AAA+ proteases, the Lon protease is known to be allosterically regulated by nucleotide and substrate binding. Although it was originally classified as a DNA binding protein, the impact of DNA binding on Lon activity is unclear. In this study, we characterize the regulation of Lon by single-stranded DNA (ssDNA) binding and serendipitously identify general activation strategies for Lon. Upon binding to ssDNA, Lon’s ATP hydrolysis rate increases due to improved nucleotide binding, leading to enhanced degradation of protein substrates, including physiologically important targets. We demonstrate that mutations in basic residues that are crucial for Lon’s DNA binding not only reduces ssDNA binding but result in charge-specific consequences on Lon activity. Introducing negative charge at these sites induces activation akin to that induced by ssDNA binding, whereas neutralizing the charge reduces Lon’s activity. Based on single molecule measurements we find that this change in activity is correlated with changes in Lon oligomerization. Our study provides insights into the complex regulation of the Lon protease driven by electrostatic contributions from either DNA binding or mutations.

**Highlights:**

- ssDNA binding allosterically activates Lon ATP hydrolysis
- Negative charge at DNA binding site is sufficient for Lon activation
- Neutralization of charge at DNA binding site inhibits Lon ATP hydrolysis
- Lon activity is linked to formation of stable Lon hexamers

**Significance:** The energy-dependent protease Lon is integral in both eukaryotic and prokaryotic physiology, contributing to protein quality control, stress management, developmental regulation, and pathogenicity. The ability to precisely regulate protein levels through targeted degradation underscores a need for tunability. We find that single-stranded DNA (ssDNA) acts as an allosteric regulator of Lon, leading to enhanced enzymatic activity. Mutations in basic residues crucial for DNA binding were found to affect Lon activity in a charge-specific manner highlighting the importance of electrostatic interactions regulating Lon’s function. Changes in Lon activity due to ssDNA binding or mutations were correlated with its oligomerization state. Our findings provide insights into the activation strategies of Lon, emphasizing the role of electrostatic contribution that modulate nucleotide affinity, oligomerization and proteolysis to advance our understanding of the complex regulatory mechanisms of the Lon protease.

## Introduction

The <u>A</u>TPase <u>A</u>ssociated with diverse cellular <u>A</u>ctivities (AAA+) family of proteases is important for regulated proteolysis in both eukaryotic and prokaryotic cells. Among this family, the Lon protease plays a particular importance in ensuring protein quality control and managing cellular stress response in bacteria (1–4). Because protein degradation is irreversible, there is a pressing need for highly controlled regulation of these proteases to ensure the rapid and specific breakdown of substates. Most proteases recognize substrates via sequence tags, known as degrons. In the case of the Lon protease, one class of these degrons are peptide motifs rich in hydrophobic residues, supporting a quality control role for the Lon protease in recognizing misfolded or unfolded proteins (5–7). Similar to all other AAA+ proteases, Lon captures the energy of ATP hydrolysis to undergo conformational changes that enable the recognition, unfolding and translocation of proteins into a nonspecific oligomeric peptidase cavity for degradation (8). In addition, Lon is particularly sensitive to allosteric regulation, with strong coupling between activities of each domain and adopting multiple distinct conformational states during the functional cycling of this protease (6,9,10).

The polypeptide product of the *E. coli lon* gene was initially identified as a DNA binding protein, originally designated CapR for its role in regulating capsular synthesis and cell elongation (11–14). The discovery of Lon as an ATP-dependent protease gave rise to considerable speculation on the role of DNA binding for the activity or function of Lon (14–21). Understanding this phenomenon was challenging because of differences in specific preparations and approaches for measuring Lon activity. For example, one study showed that denatured DNA stimulates casein degradation by Lon in an ATP dependent manner, while another demonstrated that addition of DNA limited proteolytic activity (22,23). Our recent studies revealed that in *Caulobacter crescentus* (hereafter referred as *C. crescentus*) Lon is recruited to the chromosome to clear DNA-bound proteins as a part of the genotoxic stress response, a function preserved in mitochondria (24). These varied and conflicting results led us to systematically explore the biochemical consequences of DNA binding for the Lon protease activity, using *C. crescentus* as our model system.

In this current work, we reveal that although Lon can bind both dsDNA and ssDNA, only ssDNA binding causes changes in Lon biochemical activity. Upon ssDNA binding, ATP hydrolysis of Lon increases primarily due to enhanced affinity for ATP nucleotide and protection from ADP inhibition. This heightened ATP hydrolysis subsequently leads to increased degradation of protein substrates, including known physiologically important targets of Lon. Our findings indicate that mutating basic residues that are essential for binding DNA results in loss of ssDNA binding with charge-specific consequences. Substituting these residues with negatively charged glutamates (Lon4E) induced a similar level of activation as ssDNA binding. However, neutralizing charge at this site (Lon4A) results in a poorly active Lon, compromised for ATP hydrolysis, more prone to inhibition by ADP, but still fully capable of assembling into peptidase-active Lon oligomers when substrate and nucleotide are present. We take advantage of these mutants to show that when Lon is primed to adopt an activated state, formation of the peptidase site requires only binding of ATP, but not hydrolysis, and that activation of Lon is correlated with persistent formation of higher molecular weight species.

Taken together, our results demonstrate that changes in surface electrostatics, induced by ssDNA binding or mutations at DNA binding sites, can induce formation of a state of Lon that is more readily activated for protein degradation.

## Results

### ssDNA activates Lon upon binding

Although Lon has been known to be a nucleic acid binding protein for some time, the consequences of single stranded DNA binding on Lon activity are unclear, with reports differing on whether it activates or inhibits. We took a biochemical approach to address this question as our previous work suggested that dsDNA could act as a scaffold for protein degradation (24). Interestingly, we found that while dsDNA did not affect Lon activity directly, addition of ssDNA significantly enhanced proteolysis of the model substrate casein (Figure 1A). This proteolytic activation extended to physiologically known Lon substrates, as the regulatory factors DnaA, CcrM and SciP were all degraded more rapidly in the presence of ssDNA (36 base oligonucleotide; OPC698) (Figure 1B, Supplemental Figure S1A). Using a polarization assay to monitor fluorescently labeled ssDNA binding to Lon, we found that ssDNA bound more tightly to Lon than dsDNA (Supplemental Figure S1B). Consistent with the need for ATP hydrolysis to induce degradation, ssDNA increased the intrinsic ATPase activity of Lon, whereas dsDNA did not (Figure 1C). Finally, adding ssDNA increased the peptidase activity of Lon, even without protein substrate (Figure 1D). Because oligomerization is needed to form the peptide hydrolysis site, we hypothesized that ssDNA binding may affect Lon activity through oligomerization, a direction we explore below.

**Figure 1.**
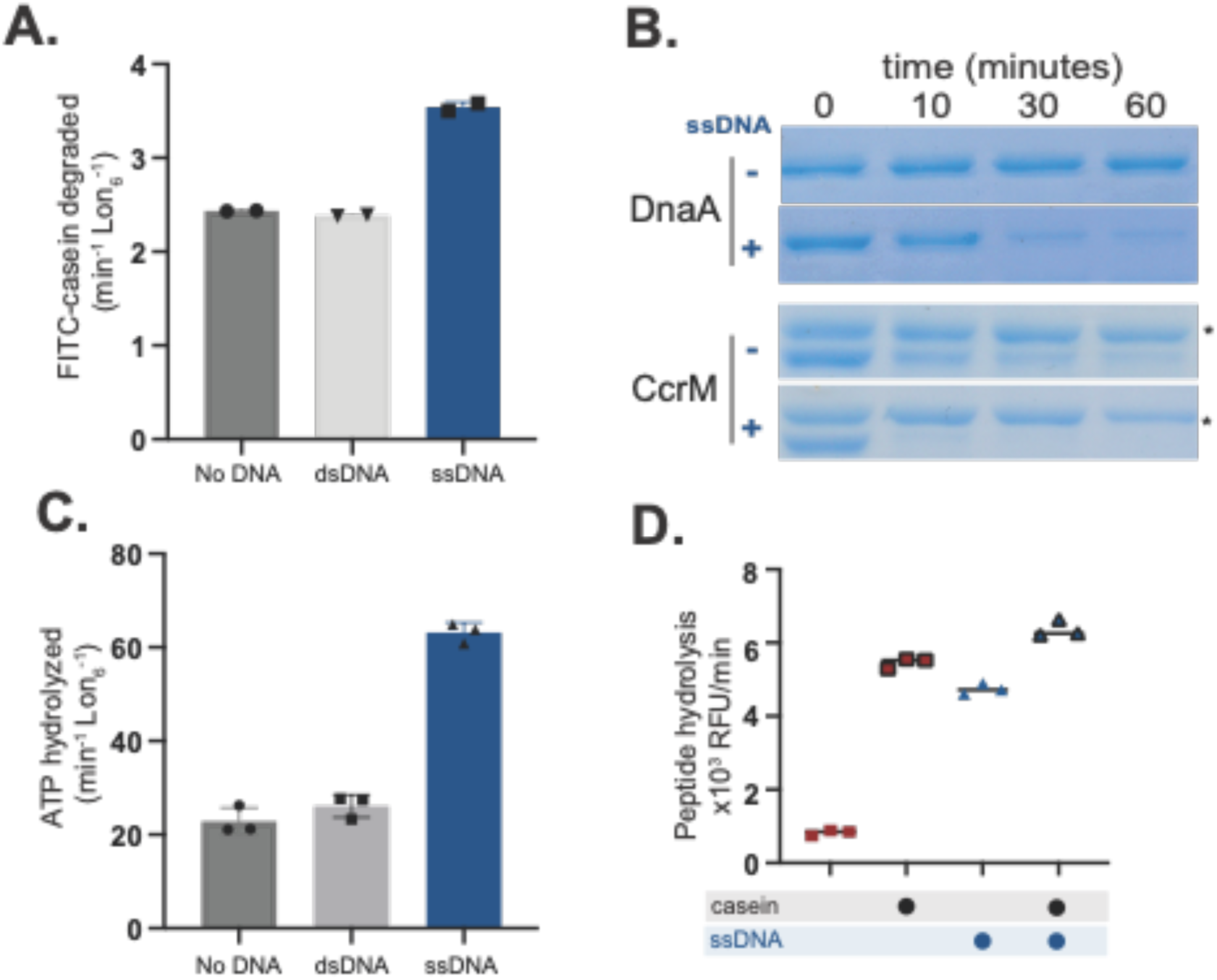
Lon binding to ssDNA increases activity. A. *In vitro* proteolysis of FITC-Casein comparing activation of purified Lon with ssDNA and dsDNA. B. Gel based *in vitro* degradation of the purified Lon substrates, DnaA (5µM) and CcrM (5µM) with and without ssDNA. Creatine kinase (*) is a component of the ATP regeneration mix. C. Basal ATPase activity with DNA using an *in vitro* ATP hydrolysis assay. D. Lon peptidase activity was obtained using the fluorogenic peptide (Glut-Ala-Ala-Phe-MNA) with and without Casein and DNA and an ATP regeneration mix with creatine kinase and creatine phosphate. All assays were performed using 0.1µM Lon_6_, 2mM ATP and 20µM DNA when noted.

As mentioned above, early investigations into the effects of DNA binding on Lon activity showed conflicting results, as some indicated that ssDNA activated Lon proteolysis (23), while others showed inhibition (22). In our initial studies, we had used various ssDNA ligands, including oligonucleotides containing G-quadruplex (G4) sequences (30 base oligonucleotide; OPC498 and 36 base oligonucleotide OJO19), a known binding motif for Lon (14,15,20). Interestingly, we observed that these G4 oligonucleotides inhibited casein degradation, but still increased stimulation of ATP hydrolysis and peptide hydrolysis (Supplemental Figure S2A, B). Additionally, like the original ssDNA described above, G4 oligonucleotides still enhanced degradation of the physiological Lon substrates DnaA and SciP (Supplemental Figure S2C). We conclude that in general ssDNA can stimulate Lon activity, but some sequences can also block degradation of a subset of substrates. We next sought to explore the basis of this activation.

### Binding of ssDNA increases Lon’s oligomerization and increases affinity for nucleotide

Lon activity is highly dependent on conformational states dictated by the presence of nucleotide and substrate. Activated Lon, bound to substrate, adopts a right-handed closed-ring spiral hexamer and inactive Lon has been found as an open-ring left-handed spiral (25). Recent structural studies further reveal the existence of intermediate oligomers that may contribute to Lon activity (26,27), supporting a general understanding that Lon adopts multiple conformational states during its activation cycle. Given that ssDNA binding directly activates Lon we hypothesized that ssDNA binding affects Lon oligomerization. To test this, we employed mass photometry to measure single particle masses of Lon alone or Lon bound to ssDNA. Our data best fits a distribution where the non-DNA bound Lon forms 75% LMW and 25% HMW. By contrast, DNA bound Lon shifts to 25% LMW and 75% higher-order active species with each Lon monomer binding to one ssDNA. This is evidenced by a MW shift from 528kDa (Lon hexamer) to 588kDa (Lon hexamer with six 11 kDa ssDNA oligonucleotides) as compared to ssDNA alone (Figure 2A, Supplemental Figure 3).

**Figure 2.**
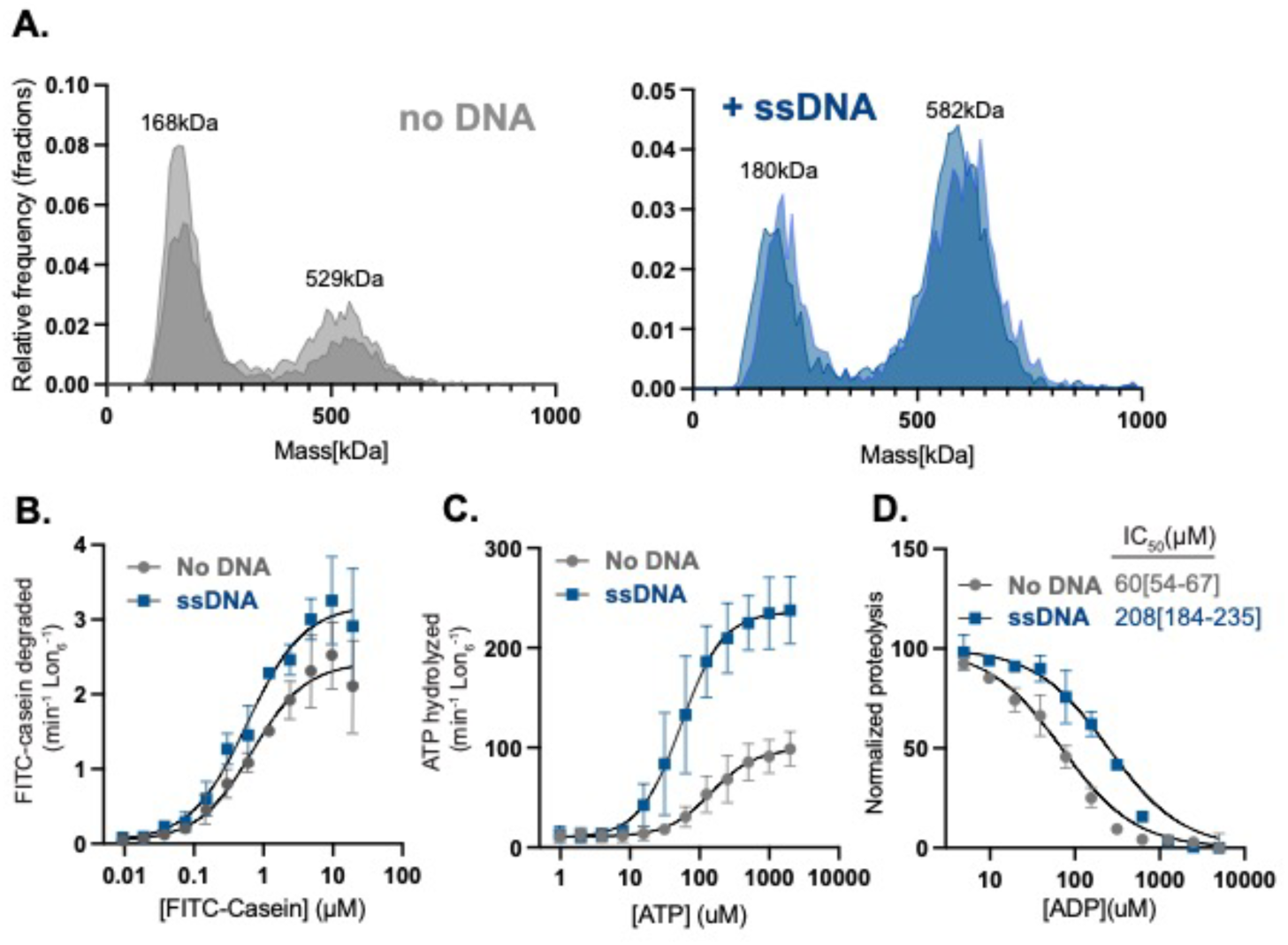
Binding of ssDNA increases oligomerization and Lon’s affinity for nucleotide. A. Mass photometry measurements of Lon with ( or without ssDNA (shaded colors represent two biological replicates). B. Michaelis-Menten plot showing the rate of degradation as a function of the concentration of FITC-Casein with and without ssDNA. C. The rate of ATP hydrolysis as a function of the concentration of ATP with 1.2µM FITC-Casein. D. FITC-Casein degradation normalized as a function of the concentration of ADP by Lon with and without ssDNA. MP experiments used 200 nM Lon monomer concentration. All other assays were performed using an ATP regeneration system with 0.1 µM Lon_6_, 20 µM ssDNA and 125 µg/mL FITC-casein with the exception or the ADP titration which did not use an ATP regeneration mix but instead used 1mM ATP to initiate the reaction.

To understand the impact of ssDNA binding on Lon activity we systematically used ATP and Michaelis-Menten kinetic experiments to determine which activities of Lon are most affected by ssDNA. First, we used saturating concentrations of ATP and titrated casein concentration while measuring initial degradation rates. We found that ssDNA increased the k_cat_ (as defined by v_max_/[Lon_6_]) but did not change the K_M_ (Figure 2B, Table 1). We next titrated ATP in the presence of saturating concentrations of casein and measured initial rates of ATP hydrolysis. Here, with the addition of ssDNA we saw an increase in the k_cat_ and a decrease in the K_M_ (Figure 2C, Table 1). Our interpretation of these results is that the primary effect of ssDNA on Lon is in modulating the interaction and/or hydrolysis of ATP. Using the Michaelis-Menten formalism with the k_on_/k_off_ representing the microscopic rate constants for ATP binding to Lon, we know that K_M_ = k_off_ / k_on_ + k_cat_ / k_on_ = K_D_ + k_cat_ / k_on_. If we assume that the on rate of ATP is only limited by diffusion, then our results suggest that at a minimum, the binding of ATP to Lon must be tighter with ssDNA present given that k_cat_ increased while K_M_ decreased with the addition of ssDNA.

**Table 1.** Steady state kinetic parameters of proteolysis and ATP hydrolysis with Lon variants and ssDNA derived from a modified allosteric sigmoidal model with the 95%CI (asymptotic) shown.

| Lon variant | Casein Degradation |  |  | ATP Hydrolysis |  |  |
| --- | --- | --- | --- | --- | --- | --- |
| | $K_M$ ($\mu\text{M}$) | $k_{\text{cat}}$ ($\text{min}^{-1}\text{Lon}^{-1}$) | Hill constant | $K_M$ ($\mu\text{M}$) | $k_{\text{cat}}$ ($\text{min}^{-1}\text{Lon}^{-1}$) | Hill constant |
| Lon | 0.7[0.6-0.9] | 2.0[1.5-2.9] | 1.1[0.5-1.7] | 92[67-135] | 88[77-103] | 1.2[0.8-1.8] |
| Lon/ssDNA | 0.6[0.3-0.9] | 3.2[2.5-3.8] | 1.1[0.5-1.6] | 54 [40-74] | 226 [201-257] | 1.4[1-2] |
| Lon4E | 1.0[0.8-1.5] | 4.0[3.6-4.3] | 1.9[1.6-2.4] | 28[23-28] | 272[246-301] | 1.6[1.2-2.3] |
| Lon4A | 0.9[0.5-1.7] | 0.2[0.1-0.2] | 2.0[1.2-4] | 206[134-707] | 45[36-75] | 1.2[0.7-1.8] |

ADP is a high affinity inhibitor of Lon even at saturating concentrations of ATP (28). We reasoned that if Lon is binding ATP tighter with ssDNA, then the Lon-ssDNA complex would be more protected from ADP inhibition. Consistent with this model, when we monitored casein degradation as a readout of Lon activity, we found that addition of ssDNA increased the IC50 for ADP 3-fold (Figure 2D). Together with the Michaelis-Menten experiments we conclude that ssDNA binding activates Lon by promoting a higher-order active species with improved ATP binding and hydrolysis, rather than altered substrate affinity.

### Mutations in the DNA binding site of Lon alter different enzymatic activities

To better understand how ssDNA impacts Lon, we explored mutations at the DNA binding sites. We previously showed a Lon variant that was unable to bind to chromosomal DNA but retained biochemical activity resulting in a physiological defect during DNA damage but not during proteotoxic stress (24). The Lon4E mutant consists of four lysine to glutamate mutations modeled from prior studies in *E. coli* (29). Our preliminary characterization found Lon4E failed to bind ssDNA (Supplemental Figure 4) but had enhanced catalytic activity, increased ATP hydrolysis and an improved ability to degrade endogenous native substrates compared to wildtype Lon (Figure 3A, B and C). This data points to either the lysine residues acting to limit Lon activity or that inversion of charge at these sites activates Lon similar to the binding of negatively charged ssDNA.

**Figure 3.**
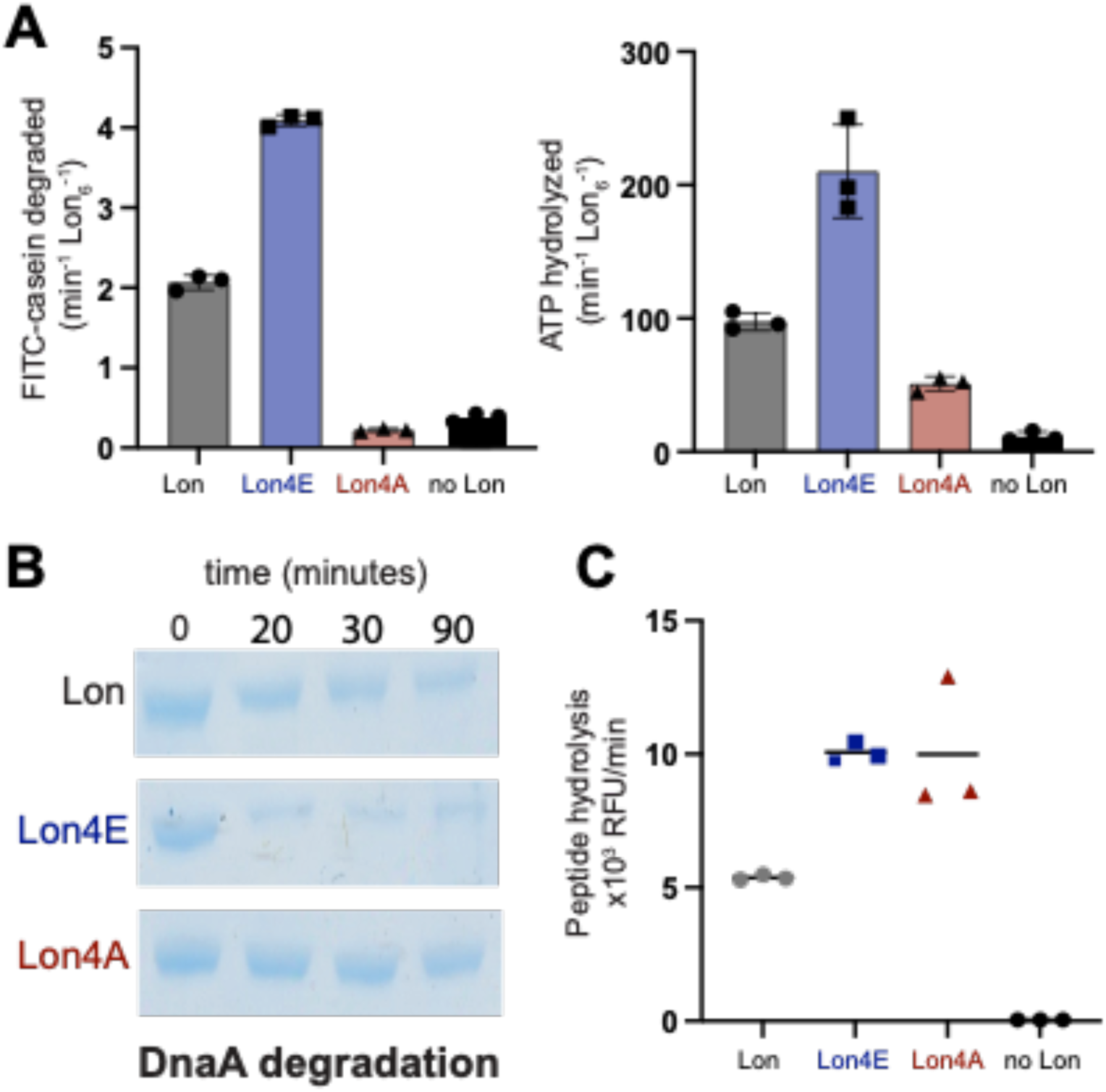
Mutations in the Lon DNA binding site alter activity. A. The following mutations K301E/A, K303E/A, K305E/A and K306E/A were introduced in the DNA binding site of Lon. Dual *in vitro* degradation and ATP hydrolysis were monitored with FITC and NADH as fluorescent readouts of proteolysis and ATPase activity. B. *In vitro* gel based degradation assay of DnaA by Lon alleles using an ATP regeneration mix and 5µM DnaA. C. Peptide hydrolysis assays with 125µM fluoro-peptide (described in Figure 1), 125ug/mL Casein and the ATP regeneration mix.

To distinguish between these two models of activation, we mutated the same lysine residues to the neutral amino acid alanine (Lon4A). We found that the Lon4A was substantially less active than wildtype for protease activity (Figure 3A and B) and ATP hydrolysis (Figure 3A). Interestingly, Lon4A retained wildtype peptidase activity in the presence of ATP with casein (Figure 3C) demonstrating that this Lon variant could still bind nucleotide and substrate to assemble the active peptide hydrolysis catalytic site. Taken together, these findings favor a model where negative charge accumulating on the surface region of Lon, either by changes of side-chain electrostatics or by binding to ssDNA, results in an allosteric activation of Lon ATP hydrolysis.

### Negative charged residues at the DNA binding site of Lon increase affinity for ATP

Based on our findings indicating that negative surface charge activates Lon we investigated whether Lon4E recapitulates ssDNA bound Lon activity by employing a series of Michaelis-Menten type experiments using the Lon variants. Negative charges at the DNA binding site induced by mutation, Lon4E, substantially increases the k_cat_ for substrate while the K_M_ remains relatively unchanged compared to wildtype Lon and the Lon4A while the Lon4A has a reduced k_cat_ (Figure 4A, Table 1). Similar to our ssDNA data, Lon4E has an increased k_cat_ with a decreased K_M_ for ATP hydrolysis, as compared to Lon4A, suggesting that a negative surface charge increases Lon affinity for ATP (Figure 4B, Table 1).

**Figure 4.**
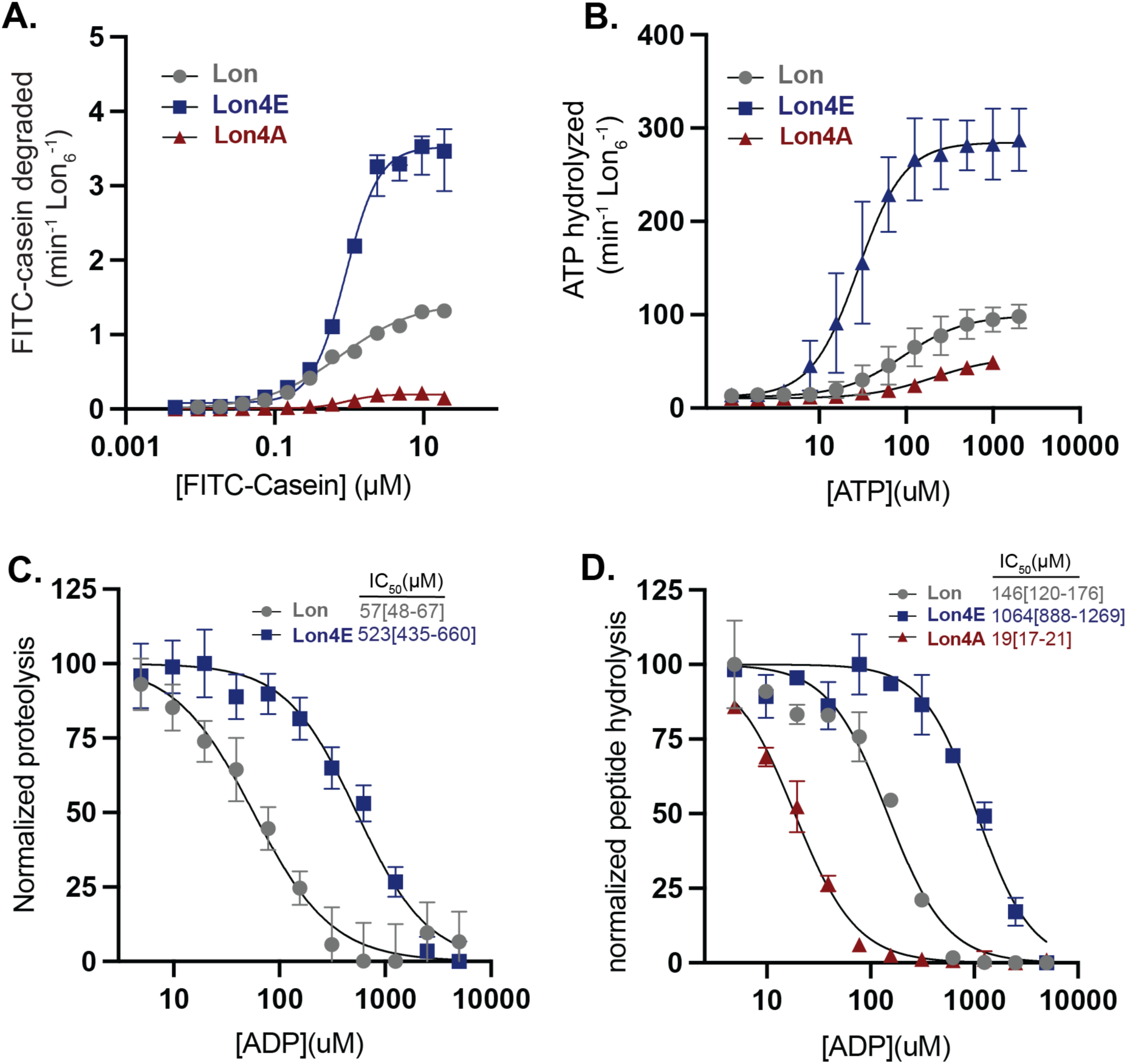
Negative surface charge at the DNA binding site of Lon increases nucleotide affinity. A. Degradation rates as a function of FITC-Casein concentration plotted using Michaelis-Menten kinetics. B. ATP hydrolysis as a function of ATP concentration. C. Normalized proteolysis as a function of ADP concentration to determine IC_50_. D. Normalized peptide hydrolysis (as described in Figure 1) as a function of ADP concentration to measure IC_50_. Experiments were performed in triplicate with the 95%CI reported.

Our model proposes that the Lon4E resembles a DNA-bound version of Lon, and if this is true we would expect a similar decrease in ADP inhibition. In support of this, we show Lon4E is less inhibited by ADP with a higher IC50 as compared to Lon for both degradation of proteins and peptidase activity (Figure 4C and D). By contrast, Lon4A, is more sensitive to ADP with a lower IC50 compared to Lon4E and wildtype Lon (Figure 4D). Together, these results strengthen our findings that a surface exposed positively charged site is allosterically coupled to ATP binding in Lon.

### Mutations at the Lon DNA binding site alter oligomerization states

To further investigate the relationship between nucleotide affinity and Lon activation we next determined how peptide hydrolysis is affected. Like for other AAA+ proteases, ATP hydrolysis is not needed for peptide bond cleavage (16,28,30,31) but formation of the peptide hydrolysis active site requires conformational changes that can be induced with nucleotide and substrate binding (25). We find that all variants of Lon could hydrolyze peptide substrates with similar rates in the presence of ATP and casein (Figure 3C, 5A and Supplemental Figure 5A). Closer examination of the traces revealed a lag phase associated with Lon4A peptidase activity (Figure 5A, Supplemental Figure 5A,C), suggestive of a need for some assembly process required prior to the activation of the peptidase active site, likely associated with the slow ATP hydrolysis of Lon4A. To separate the role of ATP hydrolysis from binding, we made use of the poorly hydrolyzed ATP analog AMP-PNP, which can stimulate peptidase activity for *E. coli* Lon (28,32). AMP-PNP alone could stimulate peptide hydrolysis with Lon4E but not wildtype Lon or Lon4A (Supplemental Figure 5B). These results suggest that the different Lon alleles may adopt different populations of conformations reflected by changes in peptide hydrolysis activity, which we sought to clarify with single molecule measurements.

**Figure 5.**
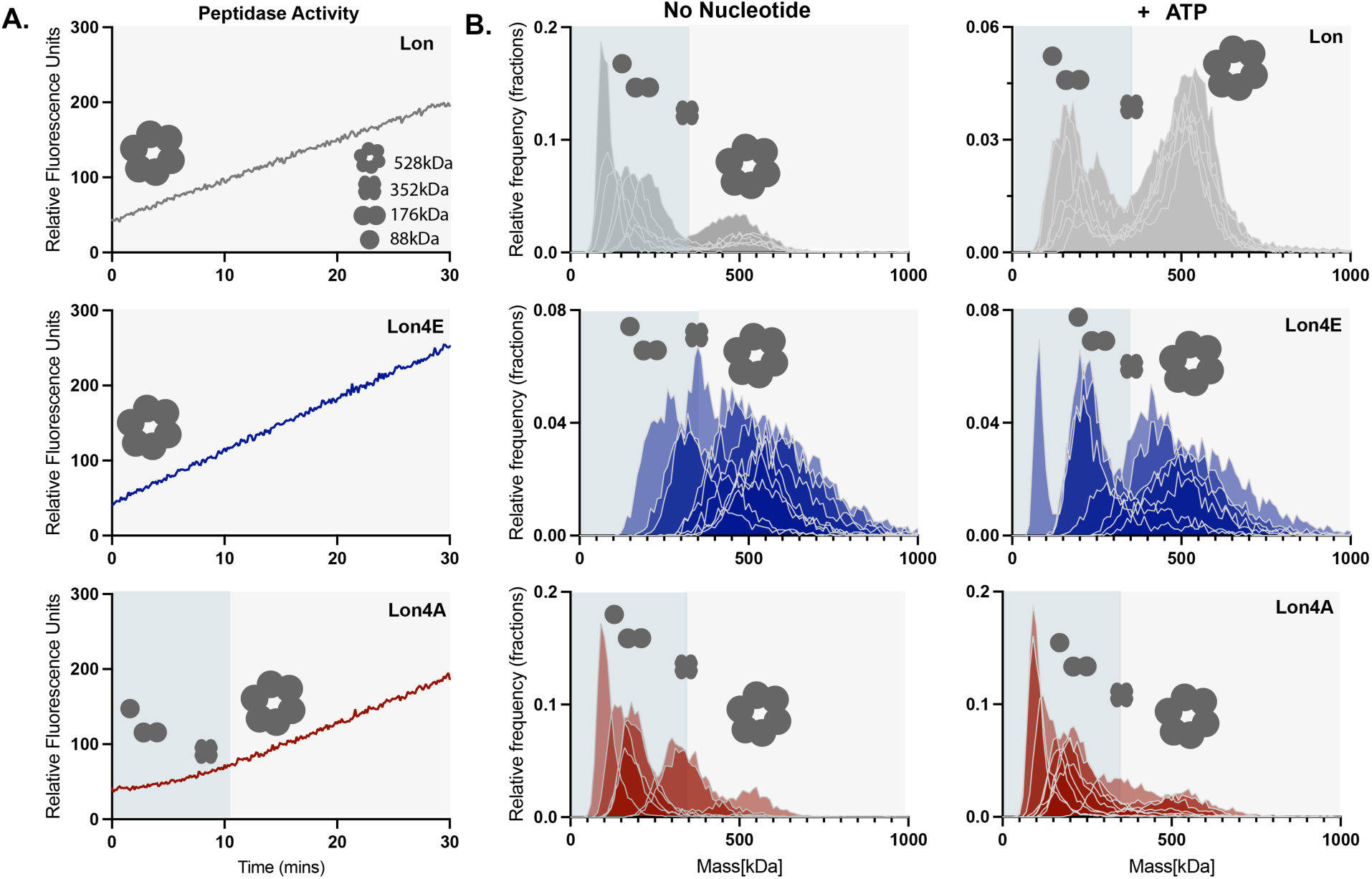
Charge state at the DNA binding site alters oligomerization. A. Peptide hydrolysis by Lon (100nM_6_) variants with an ATP regeneration mix and 125 µg/mL casein (Inset displays predicted mass of Lon oligomers). B. Mass photometry analysis of Lon variants (200 nM monomer concentration) after diluting into TK buffer and after adding saturating ATP (1 mM). Density is plotted as a relative frequency against mass (kDa) with the relative kDa for each condition. Multiple replicates for each condition (n=7) are shown.

Upon immediate dilution from concentrated stocks in the absence of nucleotide, wildtype Lon forms high molecular weight (HMW) species (>300 kDa) that are consistent with tetramer to hexamer sized complexes but primarily settles into a lower molecular species (LMW; <200 kDa) consistent with monomers to dimers (Figure 5B). By contrast, Lon4E stays in a HMW species, while the Lon4A is predominantly LMW (Figure 5B). Addition of ATP results in rapid HMW formation for wildtype Lon, but Lon4A remains in a LMW species while Lon4E maintains a HMW profile.

Our interpretation is that Lon adopts different conformations dependent on allele type which correlates with activation ability. This is particularly interesting as the sites that influence activity in this study are far removed from the oligomeric interfaces seen by structural studies, suggesting an allosteric mechanism linking these activities. The more active Lon4E persists in a dynamic range of larger oligomeric forms while Lon4A readily shifts to a monomeric/dimeric species and fails to assemble higher order oligomers as easily in the presence of nucleotide. Together, these results are consistent with stable hexameric species being the primary enzyme active state of Lon and destabilization observed when Lon cycles to an inactive state such as seen with Lon4A.

### *In vivo* characterization of Lon alleles

Finally, we tested whether these biochemical differences in Lon activity had any *in vivo* consequences under standard laboratory conditions. We generated strains carrying wildtype, Lon4E or Lon4A alleles as the sole copies of the *lon* gene. All strains grew normally under standard laboratory conditions with all of them rescuing the extended lag phase seen with *Δlon* strains (Figure 6A). Consistent with the importance of chromosomal binding of Lon in the genotoxic stress pathway (24), both Lon4A and Lon4E were sensitive to DNA damage (Figure 6A). All strains were equally resistant to proteotoxic stress generated by misincorporation of canavanine (Figure 6A). Similarly, all strains showed the same ability to degrade DnaA, a known Lon substrate, as determined by translational shut off experiments (Figure 6B). All strains were morphologically similar with respect to cell length, but both DNA-binding deficient Lon alleles result in longer stalk lengths than wildtype cells (Figure 6C, Supplemental Figure 6). Given the dramatic consequences of loss of Lon, these relatively mild effects from Lon alleles suggest that the biochemical differences we observe *in vitro* are bypassed successfully by other features important for *in vivo* activity under laboratory growth conditions, which we discuss in more detail below.

**Figure 6.**
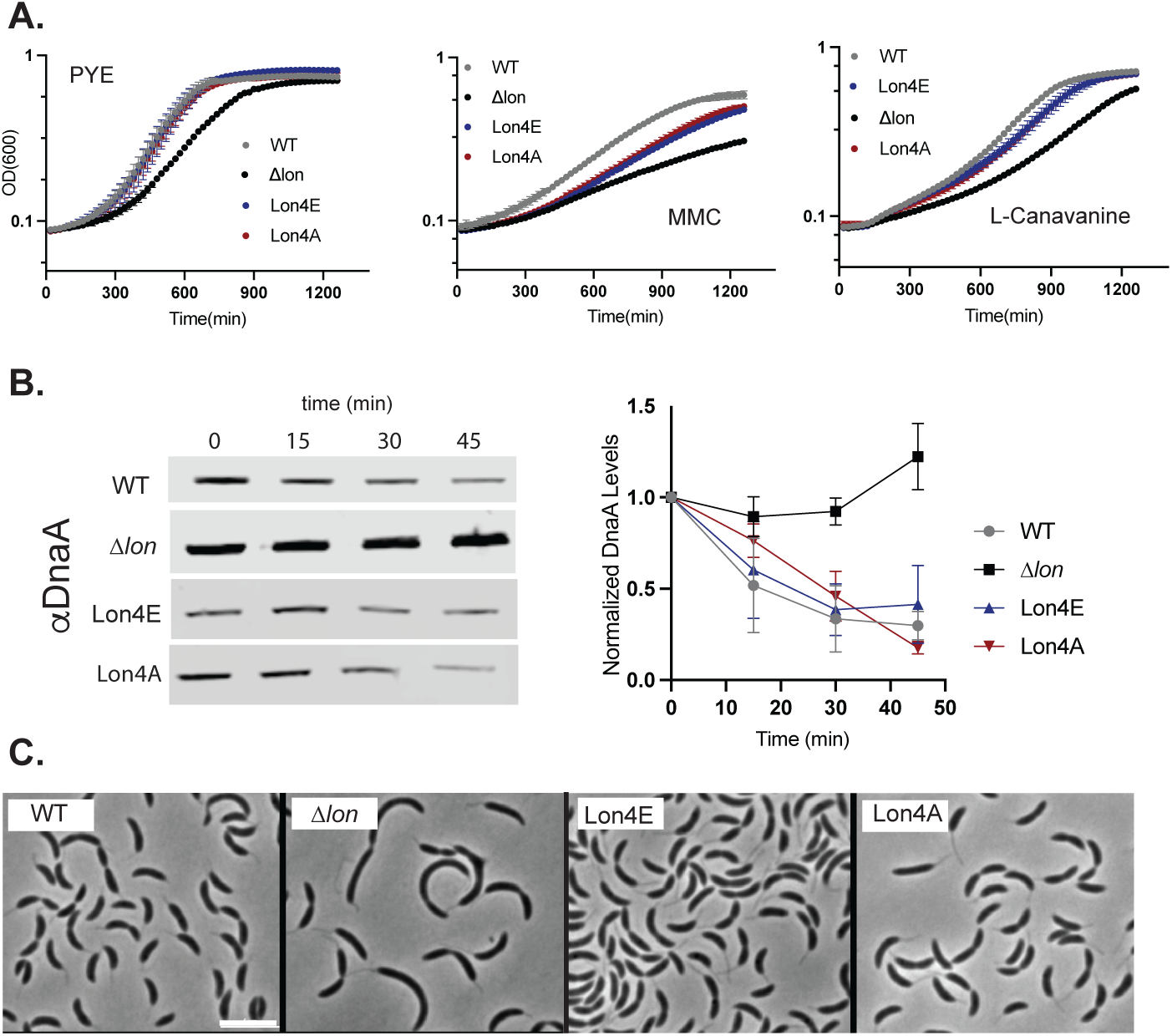
Lon alleles complement normal function with the exception of genotoxic stress. A. 24-hour growth curve of Lon variants in normal growth conditions (PYE), with genotoxic stress (0.5 µg/mL mitomycin C, MMC) and proteotoxic stress (100 µg/mL L-canavanine). B. Translational shutoff assays to monitor DnaA stability in cell. Chloramphenicol was added to stop protein synthesis and lysates were taken at the time points specified using western blot analysis with a DnaA antibody. Quantifications of six individual replicates are shown; DnaA levels are normalized to ClpP. Error bars represent SD. C. Phase contrast microscopy of exponentially growing cells, scale bar is 5 microns.

## Discussion

The Lon protease is a broadly conserved AAA+ protease found in all kingdoms of life. Lon’s activity is encoded on a single polypeptide with functional domains for protein recognition, ATP hydrolysis and peptidase activity. Mechanistically, the complex allosteric control of Lon is likely to stem in part from this linked organization of domains. Physiologically, the fact that Lon recognizes regions of high hydrophobicity rather than sequence-specific degrons allows it to function as a general protein quality protease and justifies the requirement for tight regulation to ensure that unregulated degradation does not harm the cell.

Although Lon was originally characterized as a DNA binding ATPase, the role of DNA binding for the enzymatic activity of the protease is not clear. In previous studies some reports observed that DNA can stimulate proteolysis of casein by Lon (23) while others report that DNA inhibits proteolysis by the Lon protein without affecting the ATPase activity (22). These apparent discrepancies may be explained by the specific sequences of DNA and the protease substrates used in these past studies. Our work shows that ATP hydrolysis is activated by ssDNA, but we found that G-quadruplex containing ssDNA can also inhibit degradation of unfolded artificial substrates such as casein. We interpret this to mean that although ssDNA can generally activate ATP hydrolysis, and thus degradation of native protein substrates, certain sequences also limit the ability of Lon to degrade unfolded substrates. It is tempting to speculate that this may represent the need for Lon to balance protein quality control (where misfolded protein degradation is important) with the need to degrade regulatory proteins (such as DnaA and CcrM). Additional work is needed to determine if this is true. Finally, studies in mitochondria and bacteria, including that from our own lab (24), have shown that Lon binding to DNA can facilitate degradation of DNA bound proteins. We have shown that this failure to bind DNA is particularly important during genotoxic stress (24) but may also be important for normal physiology, such as degrading the StaR repressor protein (37), which could lead to the stalk length phenotype seen in strains carrying either DNA-binding mutant alleles (Supplemental Figure 6).

Based on prior studies and our current findings, we propose some additional features of Lon activation. First, Lon binding to single-stranded DNA generally causes an increase in ATP hydrolysis due to an increased affinity of Lon for ATP. Because increased ATP consumption generally leads to increased protein substrate degradation, ssDNA-bound Lon thus degrades proteins faster, but this activation does not change the overall affinity or preference of Lon for its protein substrates. The activated form of Lon protease is less susceptible to inhibition by ADP, suggesting that either ADP binds more poorly by activated Lon or that ATP binds more tightly.

### Our kinetic data support the latter hypothesis

Second, activation of Lon can also arise from introduction of local negative charge at the DNA-binding residues of Lon (Figure 7A), with the same effects on ATP binding and substrate degradation as with ssDNA binding. Neutralization of these charges results in a Lon variant that takes more time to assemble into a peptidase active oligomer, has reduced ATP hydrolysis, and with ATP-alone, fails to readily form oligomers (Figure 7B). We conclude that electrostatic changes introduced by mutation or by ssDNA binding at specific regions that are distant from known protomer-protomer interfaces can shift oligomeric conformations of Lon; satisfying the action-at-a-distance definition of allostery (Figure 7A). Importantly, addition of ssDNA cannot act to simply tether subunits of Lon to promote oligomerization as point mutants have the same activating effects. Finally, we note that other allosteric effectors may bind to this same surface site to elicit changes in Lon activity.

**Figure 7.**
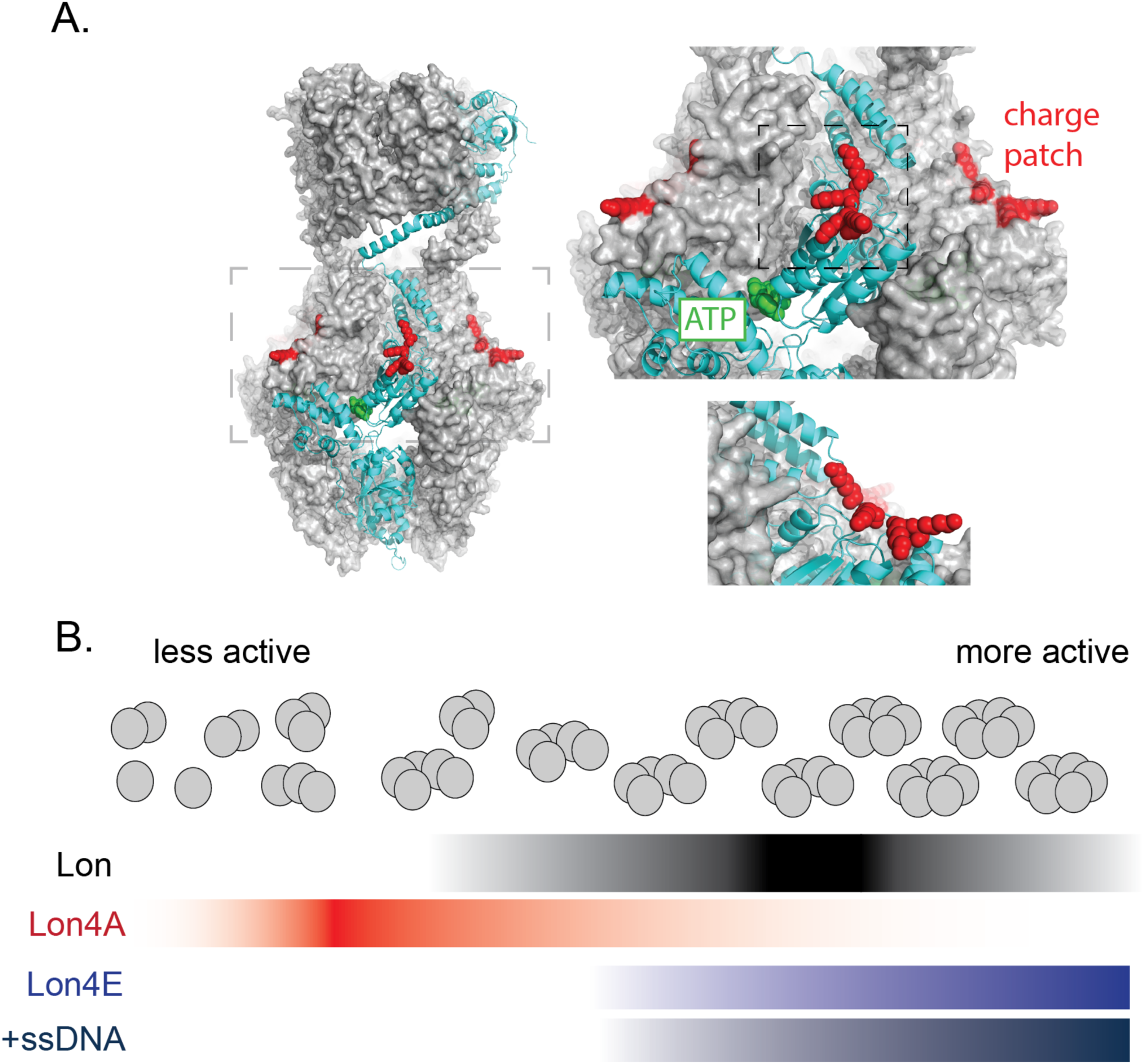
Lon activation and oligomerization are allosterically regulated by charge state at the DNA-binding sites. A. Structure of *Caulobacter* Lon as predicted by AlphaFold3. The inset displays the ATPase domain with DNA-binding residues in red (charge patch). A single monomer of the hexameric oligomer is highlighted (cyan) illustrating that the protomer-protomer interface residues responsible for ATP binding and hydrolysis (Walker A and B motif) are well-removed from the DNA-binding sites. B. Activity and oligomeric conformations of Lon depend on the overall electrostatics of the charge patch. Lon variants with negative residues at that site (Lon4E) or bound to DNA (+ssDNA) primarily form higher molecular weight oligomers. Neutralization of charge results in smaller, lower molecular weight complexes. Wildtype Lon can adopt a range of states depending on nucleotide, substrate, and effector binding.

Despite the clear biochemical differences, we do not see any substantial fitness defects or advantages when comparing strains expressing Lon variants in any laboratory conditions tested so far. One reason for this could be that crowding conditions or substrate concentrations *in vivo* are sufficiently high to promote oligomerization even with the ‘inactive’ variants. Another may be that under the growth conditions we are studying, even the 5-10% of Lon activity sustained by the ‘inactive’ variant *in vitro* may be sufficient for physiological need or stress response. Given the biochemical differences, it is tempting to speculate that other conditions may reveal more physiological impact, such as during changes in ADP/ATP ratios during nutrient starvation. Due to the importance of Lon as a quality control protease and as a cell cycle regulator it is also possible that Lon utilizes other regulatory mechanisms, such as the upregulation of the Lon activator LarA during proteotoxic stress (4) or other regulators, yet to be determined, in order to maintain proteostasis. The *in vivo* consequences of Lon activation are a fascinating topic for future studies.

Collectively, our findings point to a complex allosteric landscape of Lon that connects oligomerization state, ligand binding, local electrostatics, and enzyme activity. Given the pleiotropic impact of Lon in every system where it has been studied, we predict that accounting for this complex regulation may be important for understanding the general role of this quality control protease.

## Methods

### Cloning, Protein expression and purification

#### Protein purification

*Caulobacter* pBAD33-Lon, pBAD33-Lon4E and pBAD33-Lon4A were purified as previously described, using hydroxyapatite resin (Sigma Aldrich) and ion exchange (MonoQ) chromatography. His_6_SciP and His_6_CcrM were purified as previously described (38). DnaA was purified (39) by expressing His_6_ tagged SUMO-DnaA fusion construct and using a Ni-NTA-agarose resin, removal of the SUMO tag by Ulp1 proteolytic cleavage and a reverse Ni-NTA. Ion exchange (MonoS) chromatography using S-buffer 100mM KCl (25mM HEPES PH7.5, 200mM L-Glutamic acid potassium, 10mM MgCl_2_ and 1mM DTT) and elution buffer containing 1M KCl (25mM HEPES PH7.5, 200mM L-Glutamic acid potassium, 300mM Imidazole, 10mM MgCl_2_ and 1mM DTT).

#### Construct design and Cloning

pBAD33-Lon4A was constructed using the pBAD33-Lon as a template. 4A mutations were made via site directed mutagenesis PCR and sequence validated (Plasmidsaurus). Lon deletion strains and Lon4E cell lines were constructed as previously described (24), using the pNPTS138 plasmid. Allelic replacement of Lon4A was performed by transforming the pNPTS138-Lon4A into lon::specR cell line with primary selection on PYE with kanamycin (25ug/mL) and secondary selection on PYE with 3%(w/v) sucrose. Clonal lines were confirmed by antibiotic sensitivity and using whole genome sequencing (SeqCenter, Pittsburgh,PA).

### In vitro assays

#### Dual *In vitro* protein degradation and ATPase assay

Lon degradation and ATPase assays for all alleles, unless noted, were performed at 30°C in a Lon activity buffer [50mM TRIS pH 8.00 10mM MgCl, 100mM KCL]. Lon alleles were used at 0.1uM Lon_6_, 125ng/mL FITC-casein Type II (dissolved in water and stored at −80C) (Sigma Aldrich), 2mM ATP, 1mM phosphoenolpyruvate,10U/mL pyruvate kinase, 30U/mL lactate dehydrogenase and 0.4mM NADH (Sigma Aldrich) with or without 20µM ssDNA. Degradation and ATP hydrolysis reactions were monitored in a dual assay on a SpectraMaxM2 (Molecular Devices) in a 384 non-binding black well plate (Corning). Proteolysis was determined by an increase of fluorescence by the unquenched FITC fluorophore at the following wavelengths, Ex465nm-Em520nm. ATPase was monitored using wavelengths at Ex340nm-Em470nm by a coupled NADH-fluorescence assay where oxidation of NADH corresponds to 1:1 with ATP hydrolyzed. Rates of the reaction were determined by: FITC-casein degraded (min^-1^ Lon_6_^-1^) used the Vmax of slope/min at steady state/9.5/26/1000/[Lon_6_ allele]. ATP hydrolysis used the Vmax of slope/min at steady state/-1/2361/[Lon_6_ allele]. Degradation and ATP hydrolysis rates were fit to a modified non-linear regression model.

#### *In vitro* proteolysis assay

Each *in vitro* proteolysis assay was performed in Lon activity buffer (described in the dual in vitro assays) and an ATP regeneration mix [4mM ATP, 75 ng/mL creatine kinase and 5mM creatine phosphate] (Sigma). Lon_6_ and substrate concentrations are indicated in figure legends. Samples were preincubated at 30°C and the reactions were initiated by the addition of ATP regeneration mix. Time points taken as specified in the figure legend and concentrations were normalized with 2X SDS-loading dye and flash frozen. Samples were run on 10% (unless otherwise specified) polyacrylamide SDS-Page gels and stained with Coomassie.

#### Fluorescent Polarization assay

Purified protein was incubated at 30°C in the following buffer with 25mM Hepes pH7.5, 10 mM MgCl, 100mM KCl and 0.05% TWEEN-20. Lon alleles were used at 0.1µM (all concentrations in hexamer), 25nM DNA labeled with fluorescein (FAM) (Integrated DNA Technologies), 25nM ssDNA (OPC698) and 25nM dsDNA G1Box (24) oligonucleotides annealed (Integrated DNA Technologies). Polarization was measured on a SpectraMaxM5 microplate reader (Molecular Devices) at excitation and emission wavelengths at 460nm-540nm with 530nm cutoff.

#### Peptidase Activity

Peptidase assays were performed using Lon activity buffer, 125 nM Glutaryl-Ala-Ala-Phe-4-methyl-β-naphthylamide (Sigma-Aldrich), 0.1uM Lon_6_, 125 ng/mL casein (Thermo Fisher Scientific) and an ATP regeneration mix [4 mM ATP, 75 ng/mL creatine kinase and 5 mM creatine phosphate]. Peptide hydrolysis was evaluated as an increase in fluorescence on a SpectraMaxM2 (Molecular Devices) at excitation and emission wavelengths 335nm-410nm.

#### Mass Photometry

Mass photometry experiments were carried out using OneMP mass photometer (Refeyn LTD, Oxford, UK) at room temperature with Aquire MP software for data analysis. Experimental procedure setup was performed as previously described (40). Proteins variants were diluted to 200 nM monomeric final concentrations in 25mM TRIS PH 8.0, 10mM MgCl, 100mM KCl, 1mM TCEP and imaged immediately after dilution, 10’, 15’, and post ATP addition (1mM). Experiments were done in triplicate at different times.

### In vivo assays

#### Bacterial strains and growth conditions

All *Caulobacter crescentus* cells used in this study originated from the NA1000 strain. Liquid cultures of *Caulobacter* were grown at 30°C in a peptone yeast extract (PYE) medium containing 2g/L Peptone, 1g/L yeast, 1mM MgSO_4_ and 0.5mM CaCl_2._ Solid media conditions were grown at 30C on PYE with 1.5% bacto-agar. *E. coli* cells were grown in either liquid or solid media (1.5% agar) at 37°C in lysogeny broth (LB). Cell strains (see construct design and cloning) were cultured using antibiotics at the following concentrations: Kanamycin 50 µg/mL, Chloramphenicol 30 µg/mL, Tetracycline 15 µg/mL L, Spectinomycin 100 µg/mL.

#### *In vivo* proteolysis Assays

Protein stability *in vivo* was monitored by translational shut offs using 30 ug/mL of chloramphenicol added to exponentially growing cells (OD_600_ 0.4-0.6). At each time point specified 1mL of cells was removed, centrifuged for 5’ at 6000Xg and normalized by OD_600_ using 2X SDS loading dye and flash frozen. Pellets were boiled for 10min and centrifuged at 21000xg for 10’. Each sample was loaded onto a 10% Bis-TRIS SDS/Page gel and run at 150V for 1 hour and transferred to a nitrocellulose membrane (Cytiva). Membranes were blocked for 1 hour at room temp with 5% milk in 1X TBST (20mM TRIS and 150mM NaCl with 0.1% TWEEN-20) and primary antibodies were used with 5% milk in 1X TBST at 4°C overnight using the following dilutions:1:5000 anti-DnaA, 1:10000 anti-ClpP, 1:5000 anti-Lon. Membranes were washed 3x 5’ 1X TBST at room temp,1:15000 IRdye800 goat anti-rabbit secondary (Li-COR) in 5% milk in 1X TBST 1 hour at room temp and washed 3x 1X TBST and imaged using the Li-COR Odyssey scanner. Densitometry for degradation was determined using Fiji(41) (NIH) and plotted using Prism (Graph Pad).

#### Bacterial Characterization: Morphology, viability, and stress Assays

Cultures were diluted to OD_600_ 0.05 and grown to OD_600_ 0.5. Morphological characterization of cells was done using phase contrast microscopy (Zeiss AXIO ScopeA1). Cells were mounted onto 1% agarose-PYE pads and imaged under 100X oil immersion. Stalks were measured by Fiji(41) (NIH) and cell length was measured by MicrobeJ for ImageJ (42). Growth curves were performed by plate reader at 30C with PYE, 0.25-0.5μg/mL Mitomycin C (MMC), 100μg/mL L-Canavanine and monitored using OD_600._

#### Quantification and statistical analysis

Graphs were generated by Prism (GraphPad). Error bars represent SD n=3 and the 95%CI is reported. To determine kinetic parameters of proteolysis (r_deg_) (Eq1) and ATP hydrolysis (r_ATPase_) (Eq2) of Lon for [substrate] or [ATP] we employed non-linear regression with a hill coefficient via an allosteric sigmoidal model. Vmax is the maximum enzyme velocity, K_M_ is the Michaelis-Menten equation in the same units as [substrate], n is the hill slope =>0.

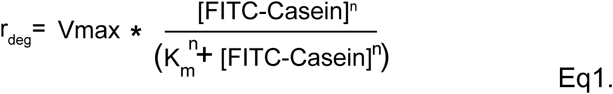

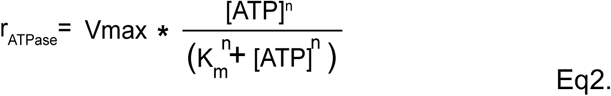

To determine the IC_50_ for ADP, data was normalized and fit using a [inhibitor] vs normalized response with variable slope model (Eq3) with the equation:

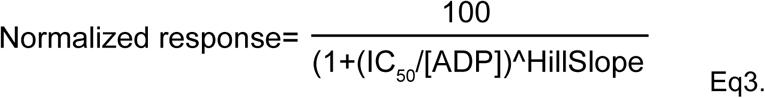

## Acknowledgments

We thank all members of the Chien lab for comments. Special thanks to Rilee Zeinert for discussion. Funding was provided for in part from NIH T32 GM235096 (JLO) and NIGMS R35 GM130320 (PC).

## Supporting Information for Ogdahl and Chien

**Supporting information Figure 1.**
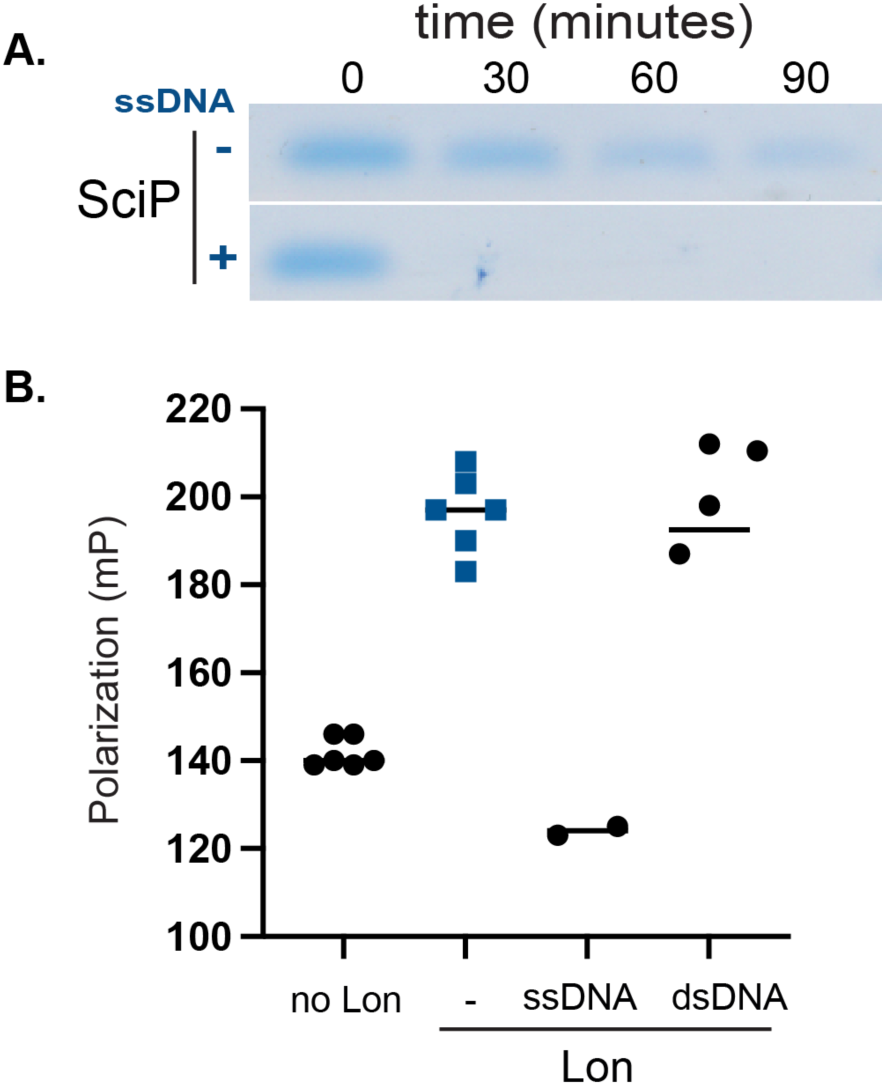
**A.** *In vitro* degradation of SciP by Lon with and without ssDNA. Assay was performed using 0.1µM Lon, 20µM ssDNA and 5 µM SciP. **B.** Fluorescent polarization assays using FAM-ssDNA and competing non-fluorescent ssDNA and dsDNA. Lon (0.1µM) measured with 25nM ssDNA.

**Supporting Information Figure 2.**
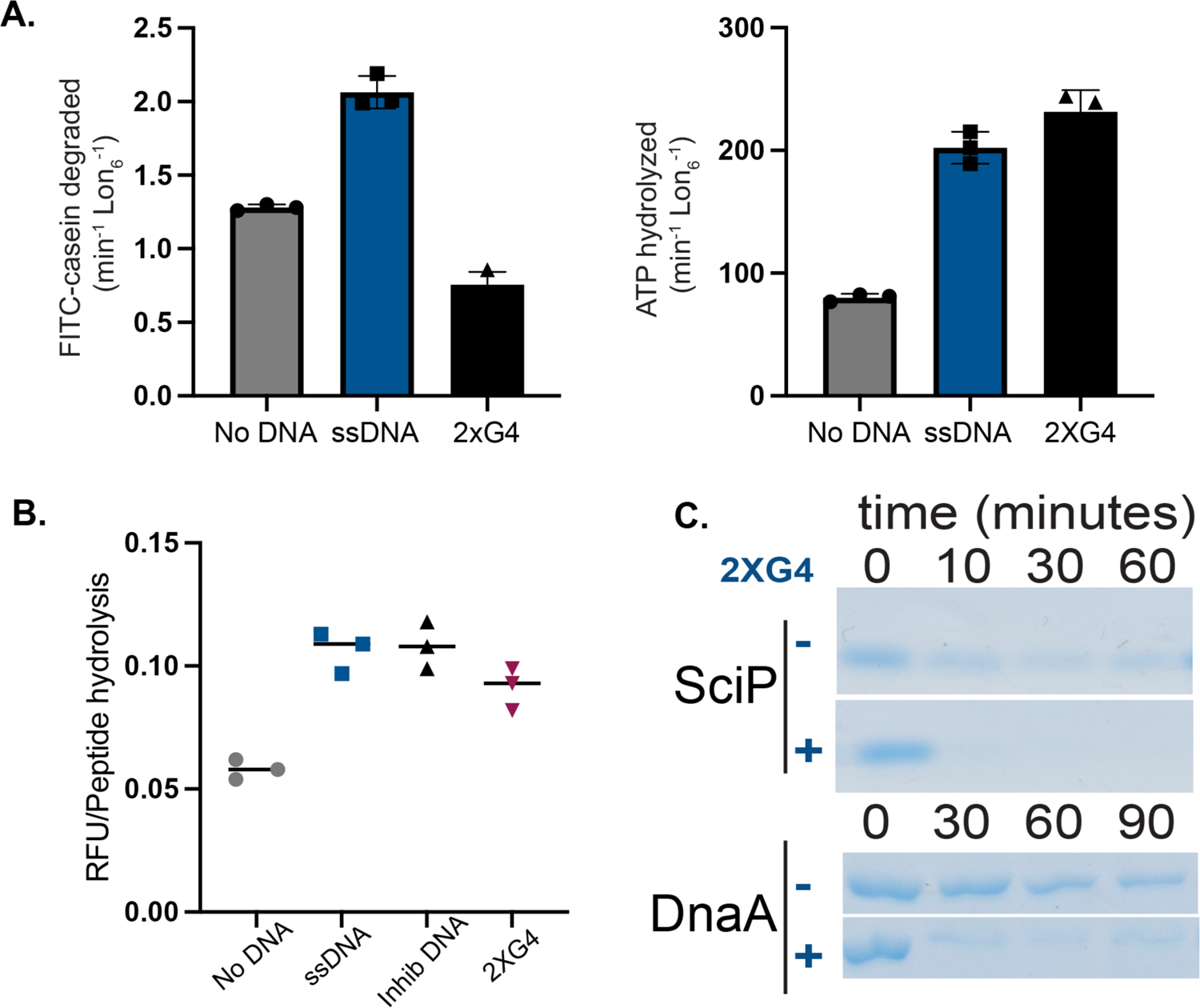
A. *In vitro* degradation and ATP hydrolysis assays using ssDNA with and without ssDNA containing 2 G4 DNA sequences (2XG4) inserted n=3. B. Peptide hydrolysis by Lon alone or with various ssDNA species (Inhib DNA contains two G quadraplexes) n=3. C. Gel based *in vitro* degradation of SciP (5µM) and DnaA (5µM) by 2XG4 ssDNA(20µM) (assay described in Figure 1).

**Supporting Information Figure 3.**
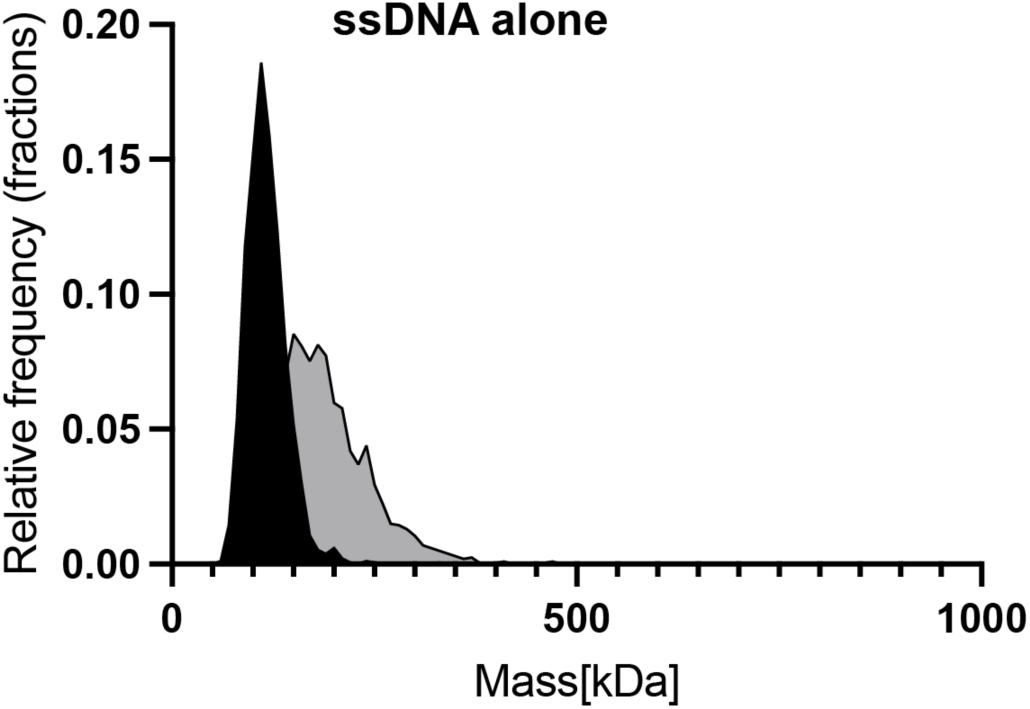
ssDNA alone does not show high molecular weight species. Mass photometry of ssDNA (20µM) alone (described in Figure 2).

**Supporting Information Figure 4.**
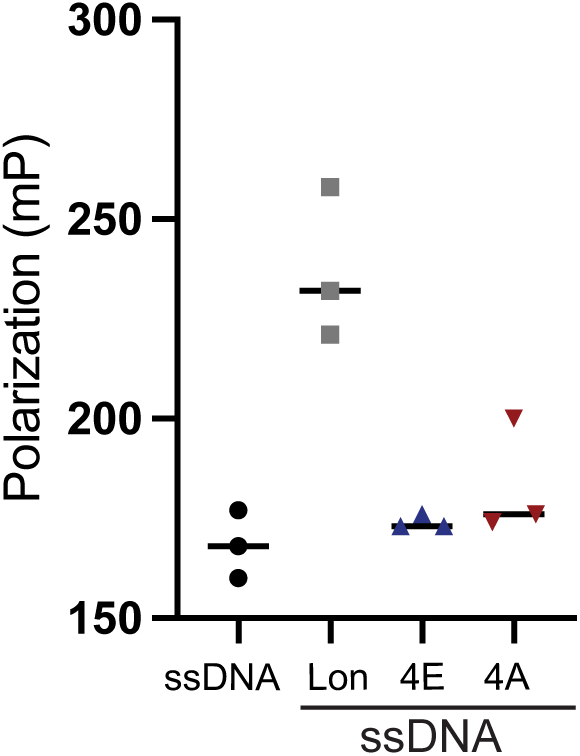
Lon4E and Lon4A do not bind to DNA. Fluorescent polarization with FAM-ssDNA and Lon variants. 0.1µM Lon variant measured with 25nM ssDNA. n=3

**Supporting Information Figure 5.**
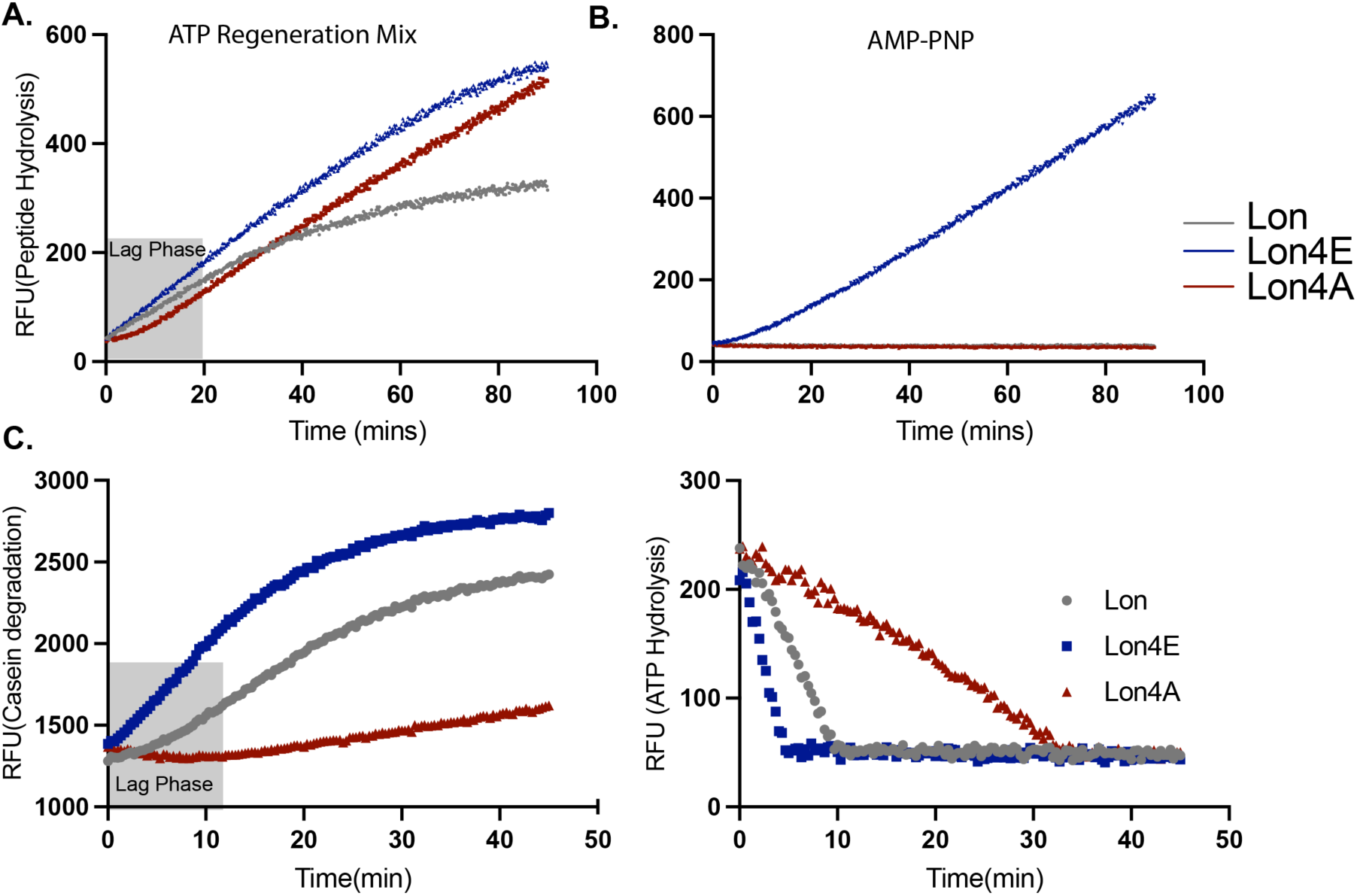
All Lon alleles hydrolyze peptides with substate and ATP, but the Lon4A has a lag period for all activity. A. Peptide hydrolysis by Lon variants of 125µM fluor-peptide (described in Figure 1) and an ATP regeneration mix, 2mM ATP, Creatine Kinase, Creatine phosphate and casein. B. Peptide hydrolysis of 125µM peptide with 1mM AMP-PNP, a slow hydrolyzing analogue of ATP, and casein. C. Raw curves of *in vitro* proteolysis and ATP hydrolysis by Lon variants. Grey box defines the lag phase for Lon4A during peptide hydrolysis and proteolysis.

**Supporting Information Figure 6.**
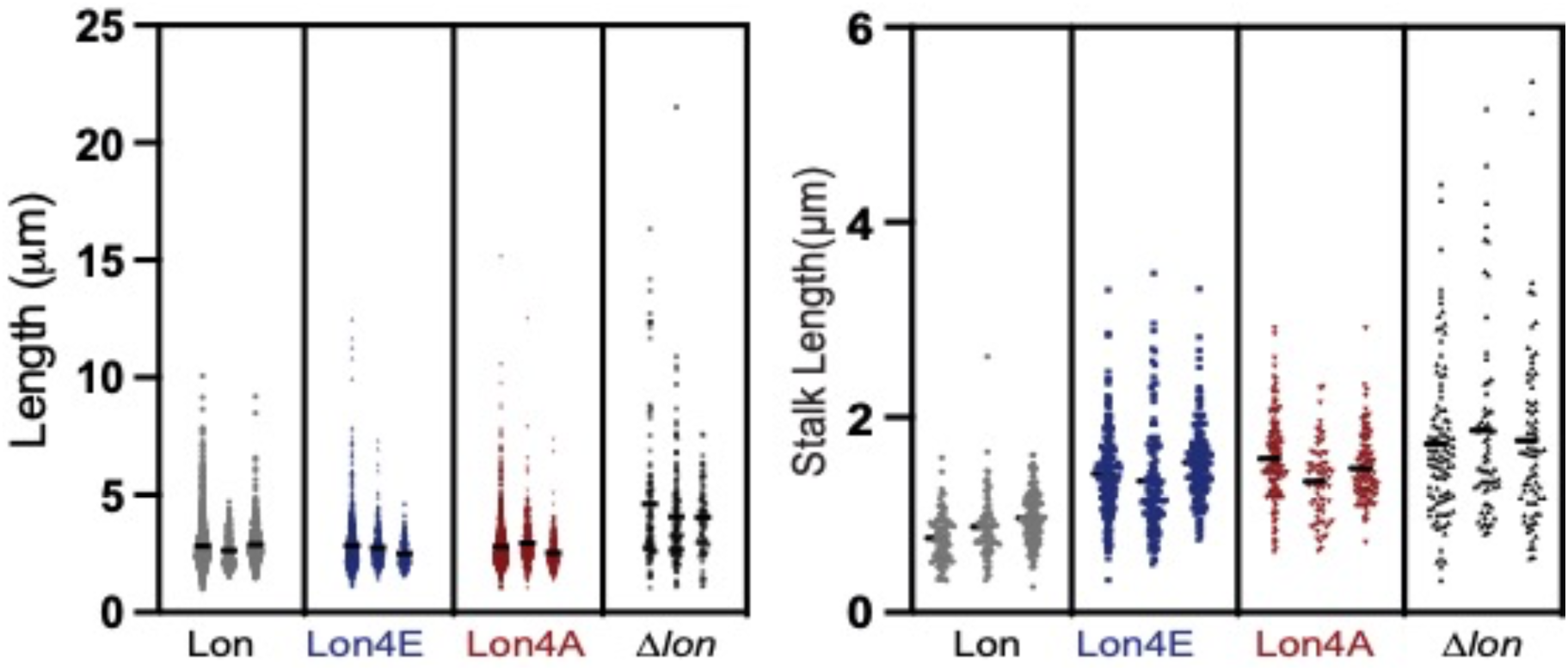
Lon4E and Lon4A have similar cell lengths but exhibit longer stalks. Cell length (MicrobeJ, Fiji) and stalk length measurements (Fiji) of the Lon alleles (representative cells shown in Figure 6) n=3.

---

**Supplemental Table 1.** Plasmids and cell strains used in this study.

| Plasmid or strain | Relevant characteristics | Comments | Reference |
| --- | --- | --- | --- |
| CPC176 | Wild type strain | NA1000 |  |
| CPC667 | $\Delta lon$ | Clean delete | Lab collection |
| CPC741 | Lon4E | Allele replacement pNPTS-138 plasmid | (24) |
| CPC1278 | Lon4A | Allele replacement pNPTS-138 plasmid for Lon4A strain | This study |
| CPC605 | lon::specR |  | (43) |
| EPC517 | pET23b-His<sub>6</sub>sumo-DnaA | BL21(DE3) Carb | (7) |
| EPC446 | pET23b-His<sub>6</sub>sumo-CcrM | BI21(DE3) Ampicillin | Lab collection |
| EPC565 | 375 SciP | BI21(DE3) Ampicillin | Lab collection |
| EPC460 | pBAD33-Lon | BL21(DE3) Chloramphenicol | Lab collection |
| EPC1504 | pBAD33-Lon4E | Top10 Chloramphenicol | (24) |
| EPC1796 | pBAD33-Lon4A | BL21(DE3) Chloramphenicol | This study |

**Supporting Information Table 2.**
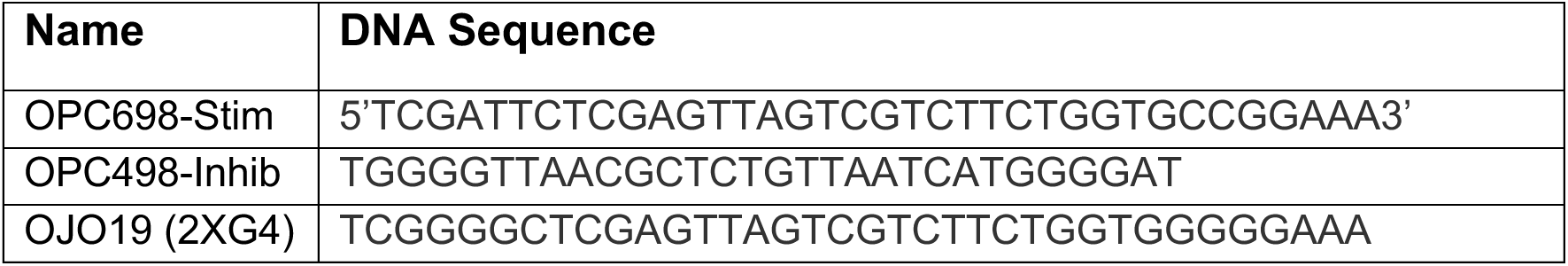
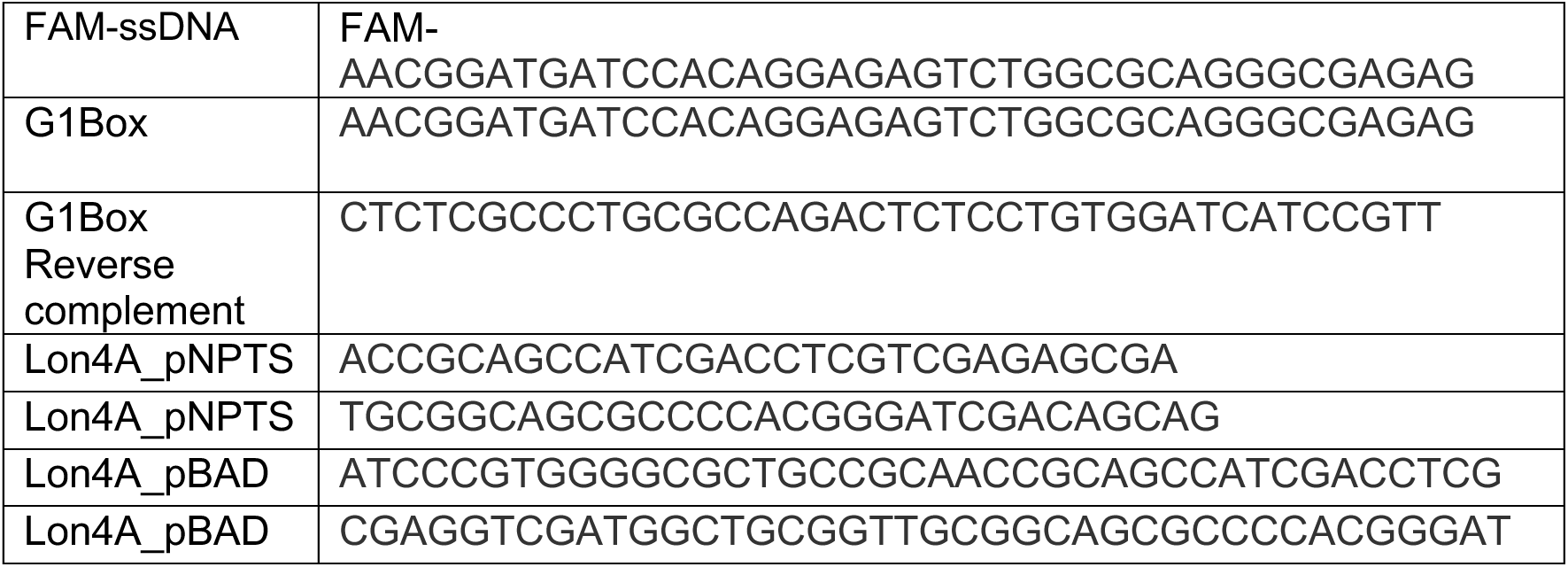
DNA sequences used in this study.

